# Exploring the Neoantigen burden in Breast Carcinoma Patients

**DOI:** 10.1101/2022.03.03.482669

**Authors:** Sambhavi Animesh, Xi Ren, Omer An, Kaijing Chen, Soo Chin Lee, Henry Yang, Melissa J Fullwood

## Abstract

In this study we performed a multi-omics analysis comprising whole-exome sequencing (WES) and RNA sequencing (RNA-Seq) on seven breast cancer patients, consisting of three Estrogen receptor (ER) positive and four Triple negative breast cancer (TNBC) subtypes to understand the neoantigen burden in breast cancer tumor samples. We predicted both class-I and class-II human leukocyte antigen (HLA) bound neoantigens by analyzing matched tumor-normal pair of exomes. Across all the patients, we predicted 434 unique neoantigens (Neo^Fil^) in total, affecting 237 different genes and 87% of them (n = 378) are expressed at RNA level (Neo^exp^). The missense mutations (87%) are the major contributor in neoantigen (Neo^exp^) generation, followed by frameshift (11%) and indels (2%). The neoantigens (Neo^Fil^) were found to be positively correlated with the somatic mutations (R^2^ = 0.89). We also noted that the vast majority (99.98%) of the predicted neoantigens are patient specific. Overall, the current study offers significant insight into the neoantigen profile in tumor types with intermediate/low mutation burdens like breast cancer.

## Background & Summary

Cancer immunotherapy has evolved into one of the most promising cancer treatment modalities. The clinical success of checkpoint blockade immunotherapy in cancer has revolutionized cancer care and our understanding of how the host patient’s immune system interacts with the tumor. The human tumor cells express antigenic determinants such as somatic neoantigens that can be recognized by the patient’s autologous T cells ^1^. These short peptides are presented by the human leukocyte antigen (HLA) molecules, leading to its recognition by CD8+ T lymphocytes and finally eliminating the cancer cells. On the contrary, the tumor cell can also escape immune surveillance by expressing specific immunoregulatory proteins that typically exist to prevent host autoimmunity and restrain excessive immune reactivity.

The advances in cancer immunogenomics and deep sequencing technology enabled the multi-dimensional characterization of genomic changes in cancers. The development of next generation sequencing helped us identify the tumor-specific mutations more accurately and precisely. These mutations give rise to “Neoantigens”, a class of major histocompatibility complex (MHC)-binding peptides that may be highly immunogenic because they are expressed only in tumor cells. Neoantigens generally arise from somatic nonsynonymous mutations in the expressed genes, resulting in a novel peptide presented to the MHC molecules^2–6^. It solely relies on recognizing mutant peptides as foreign and stimulating an immune response that destroys the tumor. Numerous previous studies have shown that the response to immune checkpoint inhibitors such as those targeting the CTLA-4 and PD-1/PDL1 often correlates with high tumor mutation burden (TMB) as well as higher numbers of predicted neoantigens, resulting in improved immunotherapy outcomes in melanoma and other cancers ^7–12^.

A number of the studies on neoantigens are targeted mainly on cancers with an exceptionally high mutation burden, such as melanoma ^4,5,12–14^ and lung cancer ^8^ etc. However, recently few studies have been implemented in cancer with an intermediate/low level of somatic mutations like breast cancer ^15–18^. Breast cancer typically harbors lower mutational loads than melanoma and NSCLC, averaging one mutation per Mb ^19^. Zhang et al. investigated the nature of the neoantigens generated and the effect of neoantigen-specific T cells in inhibiting or abolishing tumor growth in preclinical breast cancer models ^18^. In this paper, they identified 74, 33, and 55 HLA class I restricted neoantigens from three patient-derived xenografts, respectively. However, this work only investigated class I restricted neoantigens without analyzing class II, and also, the breast cancer samples used were patient-derived xenografts (PDXes). In another study, Wang et al. highlighted the correlation of low mutation and neoantigen burdens and fewer effector T cell infiltrate with lymph node metastasis in breast cancer by analyzing the data downloaded from the TCGA (The Cancer Genome Atlas) database ^17^. Most of the studies reported in the literature are conducted on mouse models or from previously published TCGA data.

In this study, we have analyzed patient tumor samples for profiling the neoantigen burden and tumor microenvironment. We employed an in-depth multi-omics approach comprising whole exome and transcriptome sequencing on patients’ tumor and matched normal cells to characterize the neoantigen signature. We also interrogated the neoantigen data from the patient sample to the tumor microenvironment.

We performed whole exome sequencing and RNA-Seq on seven breast cancer patients consisting of three Estrogen Receptor (ER) positive viz BC_Pt1, BC_Pt2 & BC_Pt3 and four Triple negative breast cancer (TNBC) subtypes viz BC_Pt4, BC_Pt5, BC_Pt6 & BC_Pt7. First, the tumor mutations were predicted using Mutect2 ^20^, and then both class-I and class-II human leukocyte antigen (HLA) bound neoantigens were predicted using the pVACSeq pipeline ^21^. On comparing the TMB with neoantigen load, a strong positive correlation was observed. Both Tumor Mutation Burden (TMB) and neoantigen load vary widely across breast cancer tumors, irrespective of their subtypes. Almost all the neoantigens were highly patient-specific (98.98%); each patient has a specific set of neoantigens.

## Methods

### Patient cohort

Breast cancer patient samples were obtained from the National University Cancer Institute, Singapore (NCIS) with patient consent, under Institute Review Board number DSRB 2008/00562. The clinical samples were collected by needle biopsy in a 1.5μL microtube by trained clinicians. All clinical samples were obtained from the National University Hospital Singapore and collected according to the Human Biomedical Research Act requirements. Informed consent was obtained for all clinical samples used in the study. The needle biopsy samples were flash-frozen in liquid nitrogen immediately and stored at −80°C until further use.

The detailed patient clinical characteristics are listed in Table S2.

### Cell culture

T2 cells were cultured using Roswell Park Memorial Institute (RPMI) 1640 media (Hyclone) supplemented with 20% heat inactivated Fetal Bovine Serum (FBS; Hyclone) and 1% penicillin/streptomycin (Hyclone) at 37°C in a CO_2_ incubator.

### Tissue procurement and nucleic acid isolation

The biopsy tumor sample and patient matched blood were collected from seven breast cancer patients. All tumor samples were flash-frozen after biopsy and stored at −80°C for DNA and RNA isolation. The blood was collected using Vacutainer CPT Cell Preparation Tube. Tumor genomic DNA and RNA were extracted from frozen tumor tissues using an AllPrep DNA/RNA Mini Kit (Qiagen). Peripheral blood mononuclear cells (PBMC) were isolated from the freshly withdrawn blood samples, and subsequently, the germline DNA was extracted using the QIAamp DNA Mini Kit (Qiagen) kit. DNA and RNA were then quantitated by the Qubit Fluorometer (Life Technologies). The RNA integrity was assessed using Agilent Eukaryotic Total RNA 6000 assay (Agilent Technologies).

### DNA whole-exome and RNA sequencing

For each patient, tumor/patient matched germline DNA samples were processed for exome sequencing. Whole-exome libraries were prepared using a SureSelectXT Human All Exon V5 kit (Agilent Technologies). The RNA seq library from tumor RNA was prepared using Illumina TruSeq Stranded mRNA Library Prep Kit as per the manufacturer’s instruction. The sequencing data were generated as 100-bp paired-end reads on a HiSeq2500 Sequencer/ HiSeq4000 Sequencer (Illumina).

### HLA Typing

The HLA subtype at four digit resolution was predicted using the seq2HLA software^22^. The input for the pipeline was standard RNA-Seq sequencing data in FASTQ format. All RNA-Seq reads were mapped to the reference HLA sequences using Bowtie ^23^. To estimate loci expression, the reads were proportionally assigned to the determined HLA loci based on the mappings between a read and the determined HLA groups. They were then normalized according to reads per kilobase of exon model per million mapped reads (RPKM)^24^. Furthermore, a confidence score was assigned to each HLA type, which reflects the likelihood that the called group is correct versus noise. The top six HLA types with lowest confidence scores were kept.

The predicted HLA types by Seq2HLA were experimentally validated by high resolution targeted sequencing of tumor DNA at CD Genomics, USA.

### Bioinformatics analysis

The whole-exome sequencing data using genomic DNAs extracted from frozen tissue samples and patient-matched peripheral blood mononuclear cells was analysed to identify somatic mutation events by Mutect2 software^20,25^. Next, the expressed 8-12 mer peptides derived from somatic mutation events together with corresponding HLA types were given as inputs for the pVACSeq ^21^ tool. Finally, we ranked and prioritized the neoantigens using stringent criteria to obtain potential neoantigens with (1) minimal variant allele frequency (VAF) of 40%, (2) tumor coverage >= 10X and (3) MT binding score < 500nM (Table S4 and S5).

### Peptide binding assay

Peptides corresponding to mutant sequences were procured from Peptide 2.0, USA, in a lyophilized form with >95% purity. The peptides were dissolved in sterile water or 10% DMSO, depending on the sequence. The binding of the candidate peptides was assessed by measuring the induction of HLA class I molecules’ surface expression on peptide pulsed T2 cells following an established protocol ^26^ with a few modifications. The T2 cells were seeded in a 24 well plate and then treated with the peptides at 10 or 100 μM concentration. The peptide pulsed T2 cells were then incubated at 37°C in a CO2 incubator for 12 to 16h. The cells were then stained with HLA-A*0201 specific mAb conjugated to FITC for 30 min at room temperature and then analyzed by flow cytometer. T2 cells with the buffer in which peptides were dissolved were used as vehicle control. The fluorescence index (FI) was calculated using the following formula: FI = [mean fluorescence intensity (MFI)_sample_ - MFI_background_] / MFI_background_, where MFI_background_ represents the value without peptide. FI>1.5 indicated that the peptide had a high affinity for HLA-A*0201 molecules, 1.0< FI <1.5 indicated that the peptide had a moderate affinity for the HLA-A*0201 molecule, and 0.5< FI <1.0 indicated that the peptide had low affinity for the HLA-A*0201 molecule.

### Tumor Microenvironment estimation

The CIBERSORT algorithm was used to predict the TIL proportions of the tumor microenvironment from tumor RNA expression profiles ^27^.

## Data Records

The datasets of RNASeq are available in GEO under accession number GSE189191 and WES are available in SRA under accession number SRR17053366

## Technical validation

### Assessment RNA integrity

Total RNA was quantified using a Qubit Fluorometer, and its quality was assessed using an Agilent 2100 Bioanalyzer according to the manufacturer’s instructions. Acceptable quality values are in the range of 1.8−2.2 for A260/A280 ratios and with an RNA integrity number (RIN) of >7.0.

### Assessment of RNA-seq data & Whole exome sequencing data quality

The quality of the raw RNA-seq data was assessed using FastQC, which ensured that the adaptors were removed from the raw reads. This program also verified that the quality of the cleaned RNA-seq reads was suitable for downstream analyses. All sequencing data were of high quality data as base call Phred quality scores above Q30, and greater than 99% of reads mapped to the reference genome.

### A comprehensive analysis of neoantigen and mutational loads in breast cancer tumor samples

To profile the neoantigen load and tumor mutational burden in breast carcinoma patients, we performed an in-depth multi-omics analysis comprising of the whole exome and RNA sequencing (Fig1). We investigated seven breast cancer patients, including three Estrogen receptors (ER) positive and four Triple negative breast cancer (TNBC) subtypes for the study. The whole-exome sequencing data using genomic DNAs extracted from frozen tissue samples and patient-matched peripheral blood mononuclear cells was analysed to identify somatic mutation events by Mutect2 software [citation]. Simultaneously, Seq2HLA was employed to predict the HLA alleles of each patient with RNA-Seq data. Finally, the expressed 8-12 mer peptides derived from somatic mutation events together with corresponding HLA types were given as inputs for the pVACSeq ^21^ tool. The peptides with binding affinities less than 200 nM were considered as potential neoantigens specific to the individual patient.

**Figure 1:**
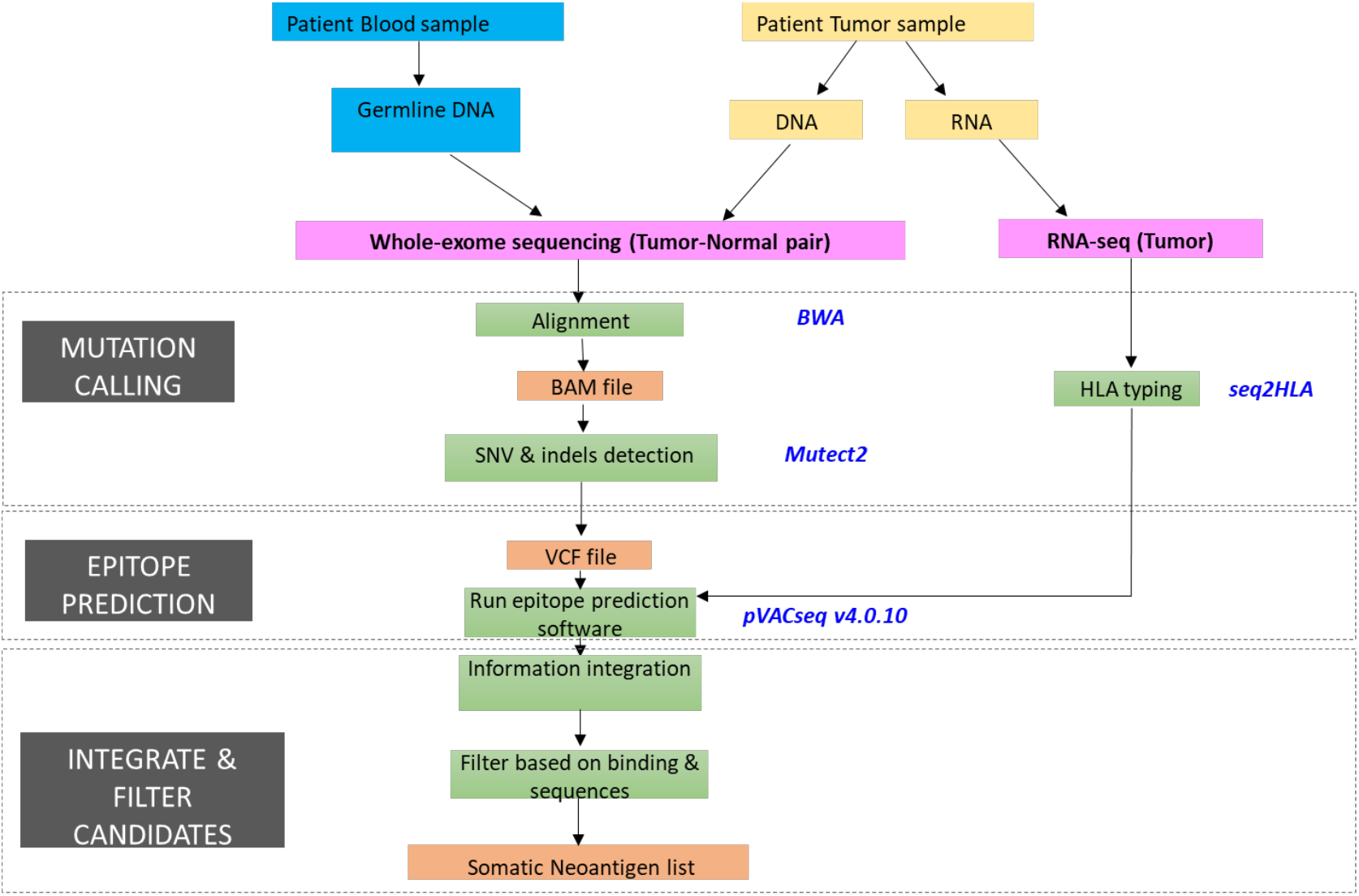
Integrative genomics and bioinformatics approach for the identification and prioritization of neoantigens in breast cancer patients. Overview of the bioinformatics strategies used for the neoantigen prediction. Whole-exome sequencing was performed on the tumor and patient matched germline DNA and RNA-Seq on tumor RNA. The somatic mutations were identified and applied in the neoantigen prediction pipeline pVACSeq (Methods).

In total, 12,136 somatic mutations and 427,347 neoantigens (Neo^ns^) (Table S1) were predicted by the software for seven patients (Fig 2C) with a median of 1390 somatic mutations and 38,622 neoantigens. The tumor mutational load and neoantigen burden of each patient was shown in Figure 2A and 2B. We found that the number of neoantigens and mutations per patient varied widely. The patient with the highest mutational load (BC_Pt7) also carried the highest neoantigen load suggesting that the TMB and neoantigen load are corelated.

**Figure 2:**
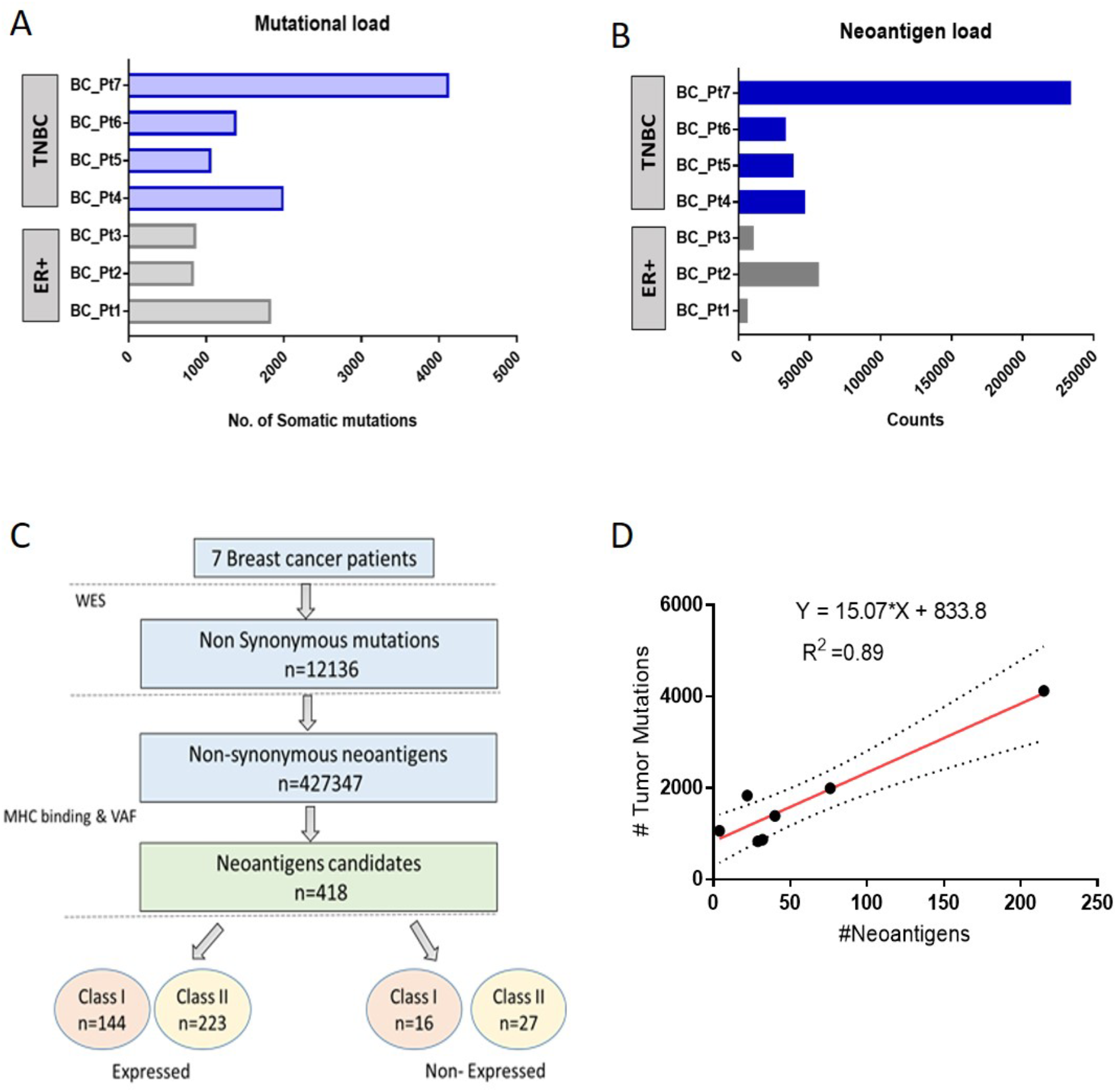
Landscape of Somatic mutations and Neoantigens profile in breast cancer patients. (A & B) The mutation burden and neoantigen load (Neo^ns^) of 7 breast cancer patients consisting of 3 Estrogen Receptor positive and 4 Triple Negative breast Cancer subtype of cancer. (C) Summary of neoantigen predictions in the 7 breast cancer patients, stratified by various filtration criteria-MHC binding, VAF(variant allele frequency), and RNA expression. (D) The correlation of the number of potential binding neoantigens (Neo^Fil^) with the number of nonsynonymous mutations calculated by Pearson correlation analysis by Graphpad prism7.03. A linear regression model is used to fit the plot’s data; the fitted line is shown in red and the 95% CIs area is shown as a dotted line around the red line.

Then, we compared the TMB with the neoantigen load of the patients. The fitting curve analysis revealed a positive linear correlation between somatic mutations and predicted neoantigens (R^2^=0.89) (Fig 2D) which were in line with the literature^8,19,28,29^ [9,20,23,24]. The prior study by Miller et al. demonstrated a positive linear relationship between mutation and neoantigen burdens (R^2^=0.862) from the MMRF CoMMpass study (NCT01454297) on 664 multiple myeloma patients^28^. Similarly, Zhou and their research group also obtained a positive linear correlation between missense somatic mutations and predicted neoantigens (R^2^=0.8845) ^29^.

Overall, our data indicate a positive correlation between TMB and neoantigen signature and the number of neoantigens increases with an increase in the number of somatic mutations in breast cancer which is in agreement with the previous published data ^8,19,28,29^.

### Comparison of neoantigens binding to the MHC class I and class II

The HLA subtyping for three major MHC class I genes (HLA-A, -B, and -C) and MHC class II genes (HLA-DP, -DQ,-DR) were called using Seq2HLA pipeline ^22^ from tumor RNASeq data and validated by targeted sequencing of genomic DNA in all the patients of our cohort. On comparing the Seq2HLA data with our targeted sequencing, we found that Seq2HLA called four-digit MHC Class I genes in all the patients with 100% accuracy. However, in MHC class II genes, the called HLA class II alleles were remarkably similar to targeted sequencing data, but we also observed slight differences (Table S3). A possible reason might be the highly polymorphic nature of the HLAII locus, where the calling in these regions is error-prone. Nevertheless, we can say that the Seq2HLA pipeline is robust and performed HLA subtyping successfully.

We then compared the fraction of neoantigens binding to MHC Class I and MHC Class II in each patient to evaluate the shared neoantigen pool between Class I and Class II molecule. We observed that 50%-75% of the neoantigens (Neo^ns^) are MHC class I specific in all the patients except in BC_Pt1, where only 38% of neoantigens are class I specific. Further, less than 10% of the neoantigens are common between Class I & II in four patients (BC_Pt2, BC_Pt5, BC_Pt6, and BC_Pt7) (Fig 3).

**Figure 3:**
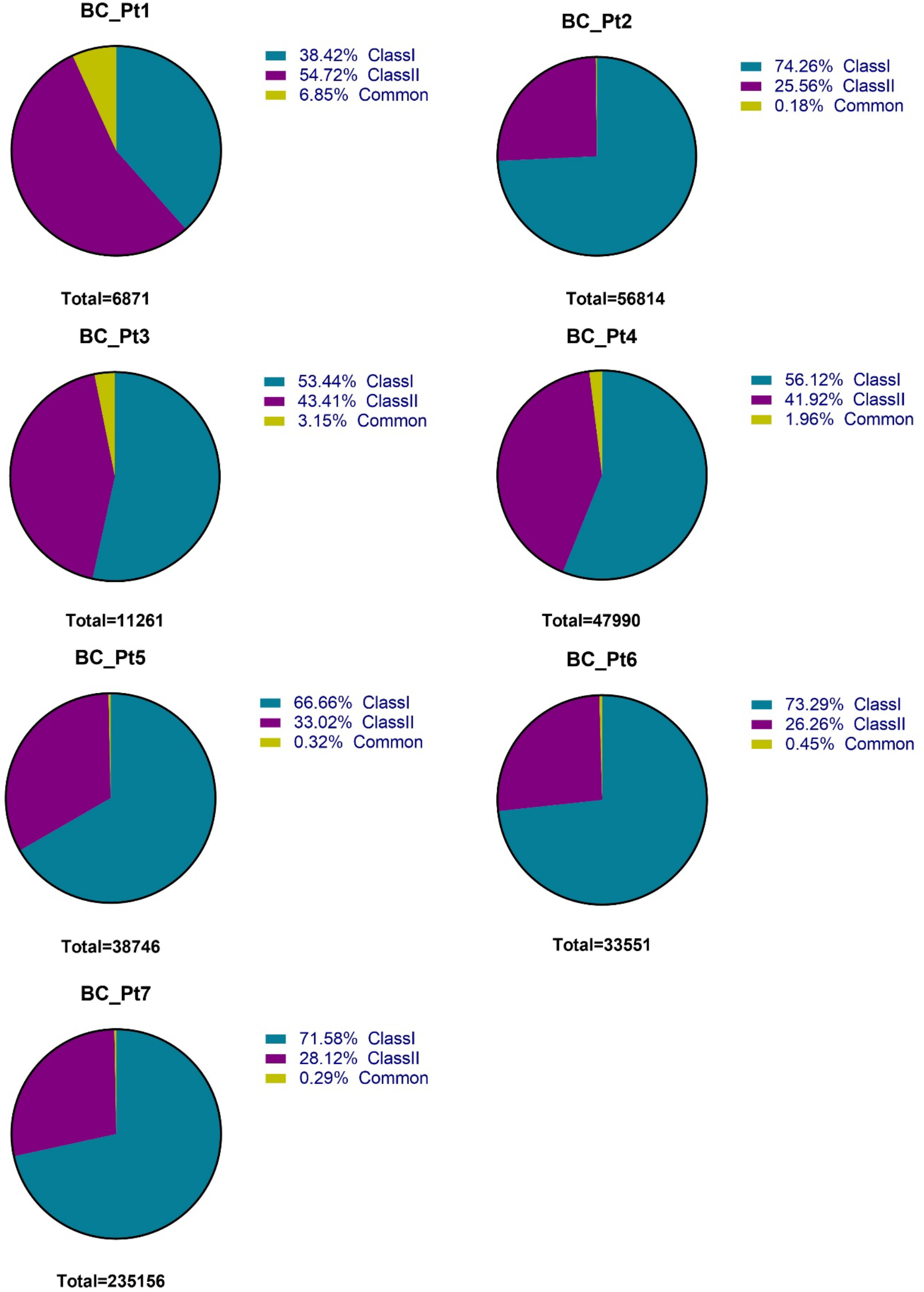
The pie chart depicting the percentage of neoantigens (Neo^ns^) binding to class I, class II and both in each patient.

Next, we sought to assess the number of neoantigens (Neo^ns^) per patient predicted to bind to each allele of HLA class I (HLA-A, HLA-B, and HLA-C) and HLA class II (HLA-DR, HLA-DQ, and HLA-DP) molecules to find the highly frequent HLA allele binding to neoantigens. We noticed that different HLA alleles bind to the different proportions of neoantigens in the patients. Amongst the class I molecule, HLA-A allele binds to the maximum number of neoantigens (Neo^ns^) in four out of seven patients (BC_Pt2, BC_Pt5, BC_Pt6 & BC_Pt7) (Fig S1A). However, among HLA DR, DQ, and DP locus of Class II MHC, HLA-DRB1 locus emerged as the strongest binder to the neoantigens (Neo^ns^) in all patients (BC_Pt1, BC_Pt2, BC_Pt3, BC_Pt5, BC_Pt6 & BC_Pt7) except BC_Pt4 (Fig S1B). The alleles HLA-C∗03:02 in BC_Pt1, HLA-A∗11:01 in BC_Pt2, HLA-B∗15:27 in BC_Pt3, HLA-C∗12:03 in BC_Pt4, HLA-A∗02:06 in BC_Pt5, HLA-A∗02:01 in BC_Pt6, and HLA-A∗33:03 in BC_Pt7 were the most frequent alleles in our cohort. Similarly, for MHC class II, HLA-DRB1∗07:01 in BC_Pt1 and BC_Pt7, HLA-DRB1∗12:02 in BC_Pt2, BC_Pt3 and BC_Pt6, HLA-DRB1∗15:01 in BC_Pt5 and HLA-DQA1∗01:02/ HLA-DQB1∗06:02 were the most frequent alleles in our study.

We also compared the neoantigens between the patients to find common, shared or recurrent neoantigens. We observed that the neoantigen profile in each patient is unique and is highly patient specific (99.98%). Moreover, the genes involved in neoantigens generations are infrequently shared. Unlike other cancers, such as gastric cancer, where researchers found six genes (PIK3CA, FAT4, BRCA2, GNAQ, LRP1B, and PREX2), which were recurrently mutated and several neoantigens were derived from these genes^29^. Zhao et al recently found that the most frequent shared neoantigens occurred in less than 2% of patients for 17 tumor types ^30^, supporting our finding that neoantigen are usually not shared among patients.

Taken together, our results suggested that the neoantigens are mostly MHC Class I specific and are rarely shared between MHC Class I and MHC Class II molecules in breast cancer. Also, HLA-A and HLA-DRB1 locus are the most frequent binder to the neoantigens in our cohort.

### Filtering and ranking strong binding neoantigens in breast cancer

The output of the pVACSeq pipeline predicted thousands of neoantigens. Consequently, we ranked and prioritized the neoantigens using stringent criteria to obtain strong HLA-binding neoantigens. The putative neoantigens were selected on applying (1) a minimal variant allele frequency (VAF) of 40%, (2) Tumor coverage >= 10X and (3) MT binding affinity score < 500nM. The lower the MT binding score the stronger the binding to MHC molecules. In addition, only the top scoring candidate per mutation was considered. As a result, we obtained 418 strong binding neoantigen candidates in seven breast cancer patients (Fig 2C).

Next, the expression of mutant alleles was examined by analysing the tumor RNA seq dataset. Any genes represented at less than 1 Transcripts Per Kilobase Million (TPM) were excluded. Consequently, we found that out of 418 neoantigens, 367 neoantigens were expressed, of which 144 were Class I specific and the rest 223 were Class II specific. The remaining 43 were not expressed (Fig 2C).

Finally, after prioritization we obtained 20, 24, 30, 66, 4, 38, and 196 candidate strong binding neoantigens (Neo^exp^) in BC_Pt1, BC_Pt2, BC_Pt3, BC_Pt4, BC_Pt5, BC_Pt6, and BC_Pt7 respectively (Fig 4A). Thus, BC_Pt5 carries the lowest neoantigen load, whereas BC_Pt7 has the highest neoantigen in our patient cohort. These neoantigens were obtained from a mutation in the expressed genes and are capable of binding to the MHC molecules.

**Figure 4:**
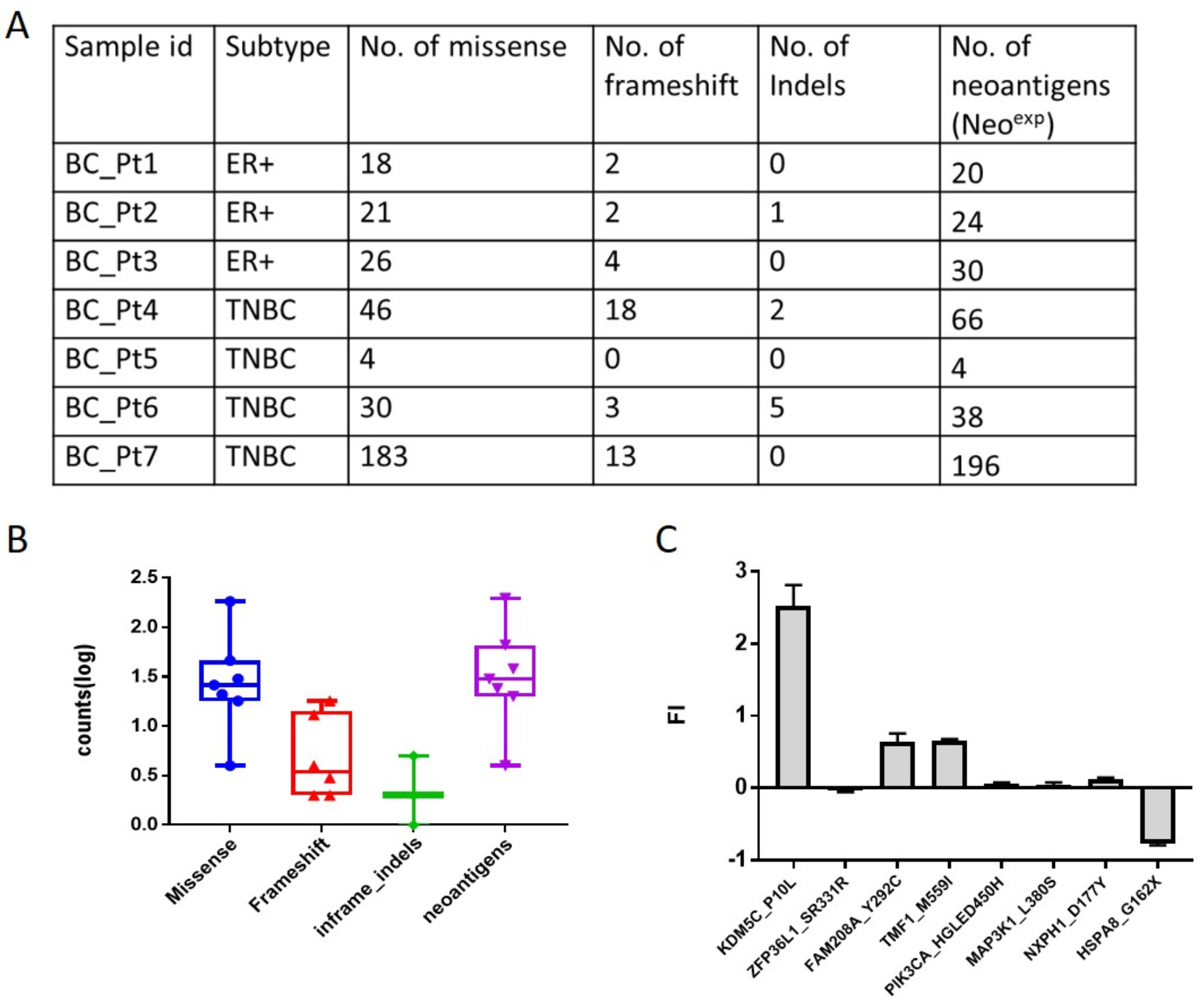
(A) The number of indels, frameshift, missense mutations and neoantigens present in each patient. (B) The boxplot shows the numbers of indels, frameshift, missense mutations, and predicted neoantigens (Neo^fil^) in 7 breast cancer patients (C) The HLA-A*02:01 specific mutant peptide binding to T2 cells

The neoantigens in our study were mainly derived either from missense, frameshift, or indels mutations. The median number is 30 for missense mutation and 3 for frameshift. Interestingly, 86.77% of Neoantigens (Neo^exp^) were derived from missense mutations as compared to frameshift (11.11%) and indels (2.11%), suggesting that missense mutations are the major contributor of neoantigen generation (Fig 4B). This finding is consistent with a previous report on gastric cancer, where most neoantigens were generated by missense mutations^29^.

Next, we wanted to check whether the candidate neoantigens can bind to the MHC molecules. To validate the binding of our candidate neoantigens, we performed a T2 cell binding assay. The T2 cells express HLA-A02:01 and are Transporter associated with antigen processing (TAP)-deficient cells; therefore, they fail to correctly translocate endogenous (processed) peptides to the site of MHC loading in the endoplasmic reticulum/Golgi apparatus [24,25]. Thus, they can be used as a model cell line to monitor the CTL response to an exogenous antigen of interest. We selected the peptide candidates for experimental validation according to the strategy described in Fig. S2. All the patients were screened for the neoantigen candidates binding to the HLA-A02:01 allotype. However, HLA-A02:01 binding neoantigens were present only in BC_Pt6. The candidate neoantigens were filtered based on their expression profile, which prioritized only eight candidates. Then, the eight HLA-A02:01 specific mutant (Mut) peptide sequences were synthesized with 99% purity (Table S6). Of the eight peptides tested, only three mutant peptides viz FAM208A_Y292C_Mut, KDM5C_P10L_Mut and TMF1_M559I_Mut bind specifically to the HLA-A02:01 allele on T2 cells. KDM5C_P10L_Mut was the strongest binder with a very high affinity for HLA-A*0201 molecules (FI-2.51). However, FAM208A_Y292C_Mut (FI-0.63) and TMF1_M559I_Mut (FI-0.66) peptide exhibit low affinity for the HLA-A*0201 molecule. On the contrary, two peptides ZFP36L1_SR331R_Mut & HSPA8_G162X_Mut failed to generate detectable peptide-specific responses (Fig 4C).

The remaining three peptides, PIK3CA_HGLED450H_Mut (FI-0.06), MAP3K1_L380S_Mut (FI-0.03), and NXPH1_D177Y_Mut (FI-0.1) bind with HLA-A*0201 molecules with almost negligible affinity (Fig 4C). It is also important to note that MAP3K1 and PIK3CA ^31–33^ are frequently mutated in breast cancer. But we observed that the neoantigens obtained from these genes do not show affinity to HLA-A*0201 molecules. Thus, our data also supports the earlier observations that not all tumor mutations can produce immunogenic neoantigens.

Finally, we observed only three candidate neoantigens out of the eight predicted binders show affinity to HLA-A02:01 in the T2 cell assay. It is interesting to note that among these eight peptides, NetMHC4 ^34^ predicted KDM5C_P10L_Mut has the highest affinity to HLA-A*0201 (MT score- 3.74), which we also observed from our experimental validation using T2 cells.

Overall, our successful detection of strong binding neoantigens in breast cancer patients highlights that the neoantigens can be predicted even in low to intermediate mutation cancers.

### The neoantigen landscape in the ER-positive and TNBC subtype of breast cancer

Next, we compared the neoantigen load between the ER-positive and TNBC subtype of breast cancer. We found that the median number of neoantigens binding to HLA molecules is more for TNBC (median = 52) than ER-positive (median = 24). However, the number of neoantigens in each patient does not depend on the breast cancer subtype. In ER-positive patients, the neoantigen varies in the range of 20-30, while in TNBC, it varies from 4-196.

The TNBC’s were often characterized with a higher mutational load and are likely to carry more neoantigens as compared to other subtypes of breast cancer ^35,36^. However, here, we observed BC_Pt7 harbors as high as 196 neoantigens (Neo^exp^), whereas patient BC_Pt5 carries only four neoantigens (Neo^exp^), although both belong to the same TNBC subtype (Fig 4A). Thus, our data points to a considerable variation in the number of neoantigens predicted (4 to 196) among the patients, indicating a significant molecular heterogeneity within breast cancer. Overall, we can say that the variation of neoantigen burden results in a high interpatient heterogeneity irrespective of the breast cancer subtype.

### The tumor microenvironment in the breast cancer

The tumor infiltrating leukocytes (TIL) status has the potential to predict cancer patients’ clinical outcomes. The tumor microenvironment with more neoantigen increases the infiltrations of TILs leading to an increase in the tumor’s antigenicity. TILs mainly include T cells, B cells, natural killer cells, macrophages, neutrophils, dendritic cells, mast cells, eosinophils, and basophils. These immune cells, found in the stroma and within the tumor itself, are often implicated in killing tumor cells.

To elucidate the immune microenvironment, we analyzed the intratumoral immune infiltrate using an in silico immunophenogram approach-CIBERSORT ^27^. The expression values of immunologically related genes were used to generate Immunophenoscores (0-10), which is a measure of the overall immunogenicity of an individual tumor, with a higher score indicative of better prognosis and better response to immunotherapy ^37^. The Immunophenoscore divided the tumor microenvironment into four aspects: antigen processing (MHC), effector cells (EC), checkpoints I immunomodulators (CP) and suppressor cells (SC). The results of tumor immune cell infiltrations showed heterogeneity across breast cancer patients. In our patient cohort, all seven patient exhibits a high score ranging from 7 to 10, indicating a very active tumor microenvironment (Fig 5).

**Figure 5:**
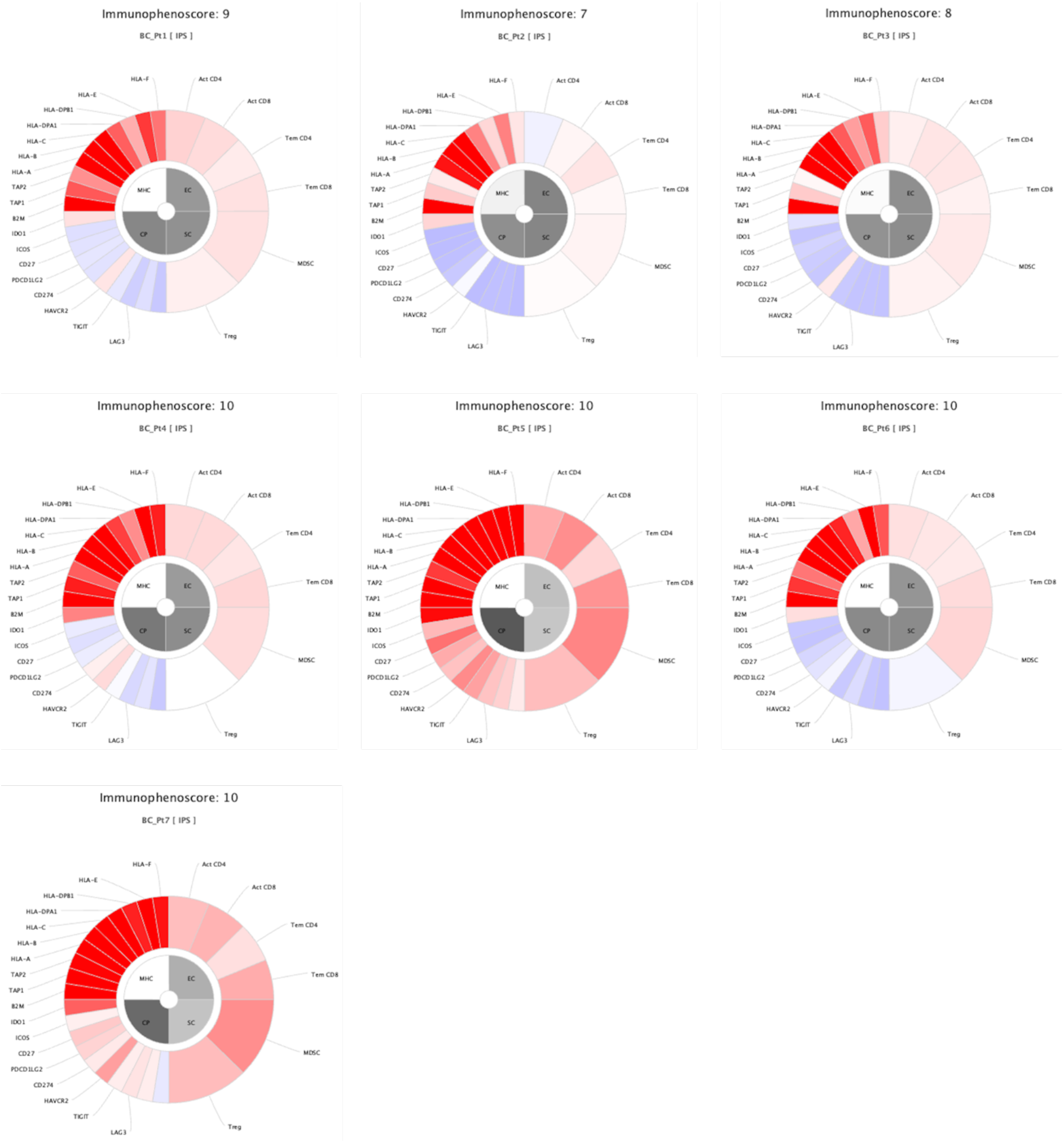
Tumor microenvironment. The expression values of immunologically related genes were used to generate Immunophenoscores (in the range of 0 to 10), which is a measure of the overall immunogenicity of an individual tumor. The Immunophenoscore of BC_Pt1, BC_Pt2, BC_Pt3, BC_Pt4, BC_Pt5, BC_Pt6 & BC_Pt7.

Previous studies have shown that the tumors with high mutational load and neoantigen load were enriched with activated T cells and were depleted with immunosuppressive Tregs and MDSCs, whereas tumors with low mutational load showed opposite enrichments and depletions^37^. In overall samples, all the seven breast cancer tumors display the infiltration of effector cells such as CD8+ T cells, activated dendritic cells. In particular, all tumors show infiltration of CD8 + T cells (6 out of 7 patients) except BC_Pt2. On the contrary, the activated Dendritic cells (DC) were observed in the tumors of only three patients (BC_Pt1, BC_Pt2, and BC_Pt6). Additionally, we also found that all tumors were also enriched with suppressor cells like M2 macrophages (Fig S3).

We also observed that BC_Pt7 has the highest neoantigen load among the seven breast cancer patients, but the tumor was enriched with suppressor cell fractions such as T regulatory cells and M2 macrophages (Fig S3). The observation of tumors with the highest neoantigen load favoring the tumor suppressor microenvironment appears to contradict earlier observations that tumor neoantigen levels are linked to immune infiltration^11,37,38^. However, McGrail et al. recently studied 19 different cancer lineages to understand determinants of CD8+ cytotoxic T lymphocyte (CTL) infiltrations in the tumors. They observed that the link of neoantigen level and immune infiltration is true across cancers characterized by recurrent mutations, but it does not hold for cancers driven by recurrent copy number alterations, such as breast and pancreatic tumors ^39^.

In conclusion, we identified breast cancer specific neoantigens, and we were able to associate neoantigen load and tumor mutation burden in breast cancer patients. We also found that missense mutation as a major contributor to neoantigen burden in breast cancer. Our data demonstrated that each patient has its own neoantigen landscape which is rarely shared. Additionally, we observed that the tumor microenvironment was not correlated with the neoantigen load in breast cancer. Thus, our data highlights the importance of assessing different tumor determinants for better monitoring of treatment efficacy. In the future, the successful design of precision vaccines for the treatment of human cancer will depend on accurate identification and prioritization of candidate neoantigens.

## Code availability

No custom script for this study. The gene expression levels were estimated by Salmon software ^40^ (https://github.com/COMBINE-lab/salmon) and neoantigen prediction was performed by publicly available pVACseq tool ^21^ (https://github.com/griffithlab/pVACtools).

## Acknowledgments

This research is supported by the National Research Foundation (NRF.) Singapore through an NRF Fellowship awarded to MJF (NRF-NRFF2012-054) and NTU start-up funds awarded to MJF. This research is supported by the RNA Biology Center at the Cancer Science Institute of Singapore, NUS, as part of funding under the Singapore Ministry of Education Academic Research Fund Tier 3 awarded to Daniel Tenen as lead PI with MJF as co-investigator (MOE2014-T3-1-006). This research is supported by a National Research Foundation Competitive Research Programme grant awarded to VT as lead PI and MJF as co-PI (NRF-CRP17-2017-02). This research is supported by the National Research Foundation Singapore and the Singapore Ministry of Education under its Research Centres of Excellence initiative. This research is supported by Ministry of Education Tier II grant awarded to MJF (T2EP30120-0020).

## References

1. Jiang, T. et al. Tumor neoantigens: From basic research to clinical applications. J. Hematol. Oncol. 12, 1–13 (2019).

2. Peng, M. et al. Neoantigen vaccine: An emerging tumor immunotherapy. Molecular Cancer vol. 18 1–14 (2019).

3. Srivastava, P. K. Neoepitopes of cancers: Looking back, looking ahead. Cancer Immunol. Res. 3, 969–977 (2015).

4. Sahin, U. et al. Personalized RNA mutanome vaccines mobilize poly-specific therapeutic immunity against cancer. Nature 547, 222–226 (2017).

5. Carreno, B. M. et al. A dendritic cell vaccine increases the breadth and diversity of melanoma neoantigen-specific T cells. Science (80-.). 348, 803–808 (2015).

6. Castle, J. C., Uduman, M., Pabla, S., Stein, R. B. & Buell, J. S. Mutation-derived neoantigens for cancer immunotherapy. Front. Immunol. 10, 1–7 (2019).

7. Spinelli, J. J. et al. Neo-antigens predicted by tumor genome meta-analysis correlate with increased patient survival. Genome Res. 743–750 (2014) doi:10.1101/gr.165985.113.

8. Rizvi, N. A. et al. Mutational landscape determines sensitivity to PD-1 blockade in non–small cell lung cancer. Science (80-.). 348, 124–128 (2015).

9. Gubin, M. M. et al. Checkpoint blockade cancer immunotherapy targets tumour-specific mutant antigens. Nature 515, 577–581 (2014).

10. Le, D. T. et al. Mismatch repair deficiency predicts response of solid tumors to PD-1 blockade. Science (80-.). 357, 409–413 (2017).

11. Van Allen, E. M. et al. Genomic correlates of response to CTLA-4 blockade in metastatic melanoma. Science (80-.). 350, 207–211 (2015).

12. Hugo, W. et al. Genomic and Transcriptomic Features of Response to Anti-PD-1 Therapy in Metastatic Melanoma. Cell 165, 35–44 (2016).

13. Gros, A. et al. Prospective identification of neoantigen-specific lymphocytes in the peripheral blood of melanoma patients. Nat. Med. 22, 433–438 (2016).

14. Lauss, M. et al. Mutational and putative neoantigen load predict clinical benefit of adoptive T cell therapy in melanoma. Nat. Commun. 8, 1–10 (2017).

15. Narang, P., Chen, M., Sharma, A. A., Anderson, K. S. & Wilson, M. A. The neoepitope landscape of breast cancer: Implications for immunotherapy. BMC Cancer 19, 1–10 (2019).

16. Ren, Y. et al. HLA class-I and class-II restricted neoantigen loads predict overall survival in breast cancer. Oncoimmunology 9, (2020).

17. Wang, Z. et al. Low mutation and neoantigen burden and fewer effector tumor infiltrating lymphocytes correlate with breast cancer metastasization to lymph nodes. Sci. Rep. 9, 1–10 (2019).

18. Zhang, X. et al. Breast Cancer Neoantigens Can Induce CD8 + T-Cell Responses and Antitumor Immunity. Cancer Immunol. Res. 5, 516–523 (2017).

19. LAlexandrov, L., Nik-Zainal, S., Wedge, D. et al. Signatures of mutational processes in human cancer. Nature 500, 415–421 (2013).

20. Cibulskis, K. et al. Sensitive detection of somatic point mutations in impure and heterogeneous cancer samples. Nat. Biotechnol. 31, 213 (2013).

21. Hundal, J. et al. pVAC-Seq: A genome-guided in silico approach to identifying tumor neoantigens. Genome Med. 2016 81 8, 1–11 (2016).

22. Boegel, S. et al. HLA typing from RNA-Seq sequence reads. Genome Med. 4, 102 (2012).

23. Langmead, B., Trapnell, C., Pop, M. & Salzberg, S. L. Ultrafast and memory-efficient alignment of short DNA sequences to the human genome. Genome Biol. 10, R25 (2009).

24. Mortazavi, A., Williams, B. A., McCue, K., Schaeffer, L. & Wold, B. Mapping and quantifying mammalian transcriptomes by RNA-Seq. Nat. Methods 5, 621–628 (2008).

25. O, A. et al. CSI NGS Portal: An Online Platform for Automated NGS Data Analysis and Sharing. Int. J. Mol. Sci. 21, (2020).

26. Hansen, T. & Myers, N. Peptide Induction of Surface Expression of Class I MHC. Curr. Protoc. Immunol. 1–8 (2004) doi:10.1002/0471142735.im1811s57.

27. Chen, B., Khodadoust, M. S., Liu, C. L., Newman, A. M. & Alizadeh, A. A. Profiling tumor infiltrating immune cells with CIBERSORT. in Methods in Molecular Biology vol. 1711 243–259 (Humana Press Inc., 2018).

28. Miller, A. et al. High somatic mutation and neoantigen burden are correlated with decreased progression-free survival in multiple myeloma. Blood Cancer J. 7, e612 (2017).

29. Zhou, J. et al. Neoantigens Derived from Recurrently Mutated Genes as Potential Immunotherapy Targets for Gastric Cancer. Biomed Res. Int. 2019, (2019).

30. Zhao, W., Wu, J., Chen, S. & Zhou, Z. Shared neoantigens: Ideal targets for off-the-shelf cancer immunotherapy. Pharmacogenomics 21, 637–645 (2020).

31. Adams, J. R. et al. Cooperation between Pik3ca and p53 mutations in mouse mammary tumor formation. Cancer Res. 71, 2706–2717 (2011).

32. Liu, P. et al. Oncogenic PIK3CA-driven mammary tumors frequently recur via PI3K pathway-dependent and PI3K pathway-independent mechanisms. Nat. Med. 17, 1116–1121 (2011).

33. Avivar-Valderas, A. et al. Functional significance of co-occurring mutations in PIK3CA and MAP3K1 in breast cancer. Oncotarget 9, 21444–21458 (2018).

34. Hoof, I. et al. NetMHCpan, a method for MHC class I binding prediction beyond humans. Immunogenetics 61, 1 (2009).

35. Liedtke, C., Bernemann, C., Kiesel, L. & Rody, A. Genomic profiling in triple-negative breast cancer. Breast Care (Basel). 8, 408–13 (2013).

36. Shah, S. P. et al. The clonal and mutational evolution spectrum of primary triple-negative breast cancers. Nature 486, 395–9 (2012).

37. Charoentong, P. et al. Pan-cancer Immunogenomic Analyses Reveal Genotype-Immunophenotype Relationships and Predictors of Response to Checkpoint Blockade. Cell Rep. 18, 248–262 (2017).

38. Tumeh, P. C. et al. PD-1 blockade induces responses by inhibiting adaptive immune resistance. Nature 515, 568–571 (2014).

39. McGrail, D. J. et al. Multi-omics analysis reveals neoantigen-independent immune cell infiltration in copy-number driven cancers. Nat. Commun. 9, 1–13 (2018).

40. Patro, R., Duggal, G., Love, M. I., Irizarry, R. A. & Kingsford, C. Salmon provides fast and bias-aware quantification of transcript expression. Nat. Methods 2017 144 14, 417–419 (2017).

